# The influence of the larval microbiome on susceptibility to Zika virus is mosquito genotype dependent

**DOI:** 10.1101/2023.05.10.540191

**Authors:** Anastasia Accoti, Laura C. Multini, Babakar Diouf, Margaret Becker, Julia Vulcan, Massamba Sylla, Dianne aster Y Yap, Kamil Khanipov, Scott C. Weaver, Mawlouth Diallo, Alioune Gaye, Laura B. Dickson

**Affiliations:** Department of Microbiology and Immunology, University of Texas Medical Branch, Galveston, Texas; Medical Zoology Unit, Institute Pasteur Dakar, Dakar, Senegal; West African Center for Emerging Infectious Diseases, Centers for Research in Emerging Infectious Diseases; Laboratory Vectors & Parasites, Department of Livestock Sciences and Techniques Sine Saloum University El Hadji Ibrahima NIASS (USSEIN), Kaffrine Campus, Senegal; Department of Pharmacology and Toxicology, University of Texas Medical Branch, Galveston, Texas; Institute for Human Infections and Immunity, University of Texas Medical Branch, Galveston, Texas; Center for Vector-borne and Zoonotic Diseases, University of Texas Medical Branch, Galveston, Texas

## Abstract

The microbiome of the mosquito *Aedes aegypti* is largely determined by the environment and influences mosquito susceptibility for arthropod-borne viruses (arboviruses). Larval interactions with different bacteria can influence adult *Ae. aegypti* replication of arboviruses, but little is known about the role that mosquito host genetics play in determining how larval-bacterial interactions shape *Ae aegypti* susceptibility to arboviruses. To address this question, we isolated single bacterial isolates and complex microbiomes from *Ae. aegypti* larvae from various field sites in Senegal. Either single bacterial isolates or complex microbiomes were added to two different genetic backgrounds of *Ae. aegypti* in a gnotobiotic larval system. Using 16S amplicon sequencing we show that similarities in bacterial community structures when given identical microbiomes between different genetic backgrounds of *Ae. aegypti* was dependent on the source microbiome, and the abundance of single bacterial taxa differed between *Ae. aegypti* genotypes. Using single bacterial isolates or the entire preserved complex microbiome, we tested the ability of specific microbiomes to drive differences in infection rates for Zika virus in different genetic backgrounds of *Ae. aegypti*. We observed that the proportion of Zika virus-infected adults was dependent on the interaction between the larval microbiome and *Ae. aegypti* host genetics. By using the larval microbiome as a component of the environment, these results demonstrate that interactions between the *Ae. aegypti* genotype and its environment can influence Zika virus infection. As *Ae. aegypti* expands and adapts to new environments under climate change, an understanding of how different genotypes interact with the same environment will be crucial for implementing arbovirus transmission control strategies.

## Introduction

Arthropod-borne viruses (arboviruses) transmitted by mosquitoes represent a major cause of morbidity and mortality worldwide [1, 2]. The mosquito, *Aedes aegypti* is the main vector for arboviruses worldwide including dengue (DENV), Zika (ZIKV), yellow fever (YFV), and chikungunya (CHIKV) viruses. Climate change and a warming world exacerbate this risk from vector borne diseases [3, 4] and arbovirus epidemics are poised to be a major threat in sub-Saharan Africa [5]. Given the ongoing and increasing risk of mosquito-borne viruses, especially in Africa, it is crucial to understand factors that contribute to their emergence and transmission.

*Aedes aegypti* demonstrates large phenotypic variability in its interactions with arboviruses largely driven by genetic and environmental variation. *Aedes aegypti* is genetically diverse worldwide [6–8] and the ability of *Ae. aegypti* to be an efficient vector of arboviruses is a partially genetically controlled trait [9]. Variation in vector competence has been widely documented among different genotypes of *Ae. aegypti* [10–12]. In some cases, vector competence is dependent on the specific mosquito-virus interactions [13–15].

Additionally, abiotic (non-living) environmental factors [16–29] and biotic (living) environmental factors [30–36] are known to contribute to the vector competence of *Ae. aegypti*. An important biotic ecological parameter influenced by the environment is the larval microbiome. Globally, *Ae. aegypti* occupies a variety of environments and diverse larval habitats. Outside Africa, domesticated *Ae. aegypti* oviposits in human artificial containers such as discarded buckets or cans and tires around human habitats. In Africa, *Ae. aegypti* will oviposit and develop in a variety of container types ranging from human artificial containers such as discarded buckets, cans, or tires around human habitats, to tree holes and rockpools in forested habitats. Larval development sites represent different microbiomes [37]. The larval microbiome is largely determined by the environment and is critical for establishing the nutritional status of the mosquito [38]. Importantly, interactions between *Ae. aegypti* larvae and different bacterial isolates drive variation in DENV susceptibility in adults [37, 38]. However, whether different genotypes of *Ae. aegypti* interact differently with the same larval microbiome to drive variation in arbovirus susceptibility is remains unknown.

Here we expand on previous work [37] to determine if the impact of the larval microbiome on arboviruses susceptibility is dependent on mosquito genotype. Using single bacterial isolates, we demonstrate that the influence of specific bacterial isolates on adults ZIKV susceptibility is dependent on mosquito genotype. Using complex microbiomes isolated from larvae in Senegal, we demonstrate that different genotypes of *Ae. aegypti* retain different members of the larval microbiome and that ZIKV susceptibility is dependent on the specific pairing between *Ae. aegypti* genotype and complex microbiome. Our results provide empirical evidence that *Ae. aegypti* genotype by environment interactions drive variation in ZIKV susceptibility. As the earth becomes warmer and drier and de-forestation and urbanization increase, *Ae. aegypti* will expand into new environments [39, 40] and exploit different oviposition container types. This is especially true in Africa where the larval development site is tied to the environment. *Aedes aegypti* genotypes accustomed to ovipositing in forest or natural breeding sites will be forced to adapt and oviposit in urban artificial container types. Understanding how climate driven changes to the environment will increasingly shape the ability of *Ae. aegypti* to transmit important human pathogens is required to guide the development and deployment of control strategies.

## Results

### Single bacterial Isolates

Previously we observed that larval development in the presence of different single isolates alters DENV replication in *Ae. aegypti* [37], but it remains unknown if the influence of larval development in the presence of different single bacterial isolates on arbovirus infection is consistent across *Ae. aegypti* genotypes. To address this, we isolated bacteria colonies from larvae collected in Thiés, Senegal. A total of 64 bacteria were isolated from larvae homogenized in 1X PBS originating from a collection site in Thies. Three bacteria isolates were chosen (*Serratia spp.* (Bacterial Isolate A), *Chyrseobacterim spp.* (Bacterial Isolate B), and *Serratia spp.* (Bacterial Isolate C)) for further characterization (see Material and Methods for information on bacteria selection). To understand if larval development in the presence of specific bacterial isolates alters pupation rates in an *Ae. aegypti* genotype dependent manner, two lines of *Ae. aegypti* from Senegal with known genetic differences [7], Thiés (THI) and Kedougou (KED), were exposed to equal amounts of bacteria isolates in gnotobiotic system and the proportion of pupae was recorded each day. Pupation rates were measured in triplicate gnotobiotic flask daily from the onset of pupation (day five post hatching) until day 10. Overall, the THI line of *Ae. aegypti* pupated slower than the KED line (p. value < 0.0001). In both lines of *Ae. aegypti*, the time it took to reach 50% pupae (PD_50_) was different between all three isolates (Figure 1A) (Supplemental Table 1). In both lines the bacterial isolate that had the slowest pupation rate was Bacterial Isolate A with a PD_50_ of 8.5 days in the THI line and 7.4 days in KED line. Conversely, the bacterial isolated that resulted in quickest pupation rate was Bacterial Isolate B where it took 6.9 days in the THI line and 6.1 days in the KED line. Bacteria C resulted in intermediate pupation rate of 7.4 days in the THI line and 6.7 days in the KED line. Interestingly, Bacteria A and C both belong to the genus *Serratia* but exert different effects on pupation rates. Overall, we observed that the PD_50_ was dependent on the mosquito genotype, bacterial isolate, or the interaction between the two variables (ANOVA on a general linearized model: mosquito genotype: p < 0.0001, bacterial isolate: p < 0.0001, mosquito genotype x bacterial isolate: p < 0.0001).

**Figure 1:**
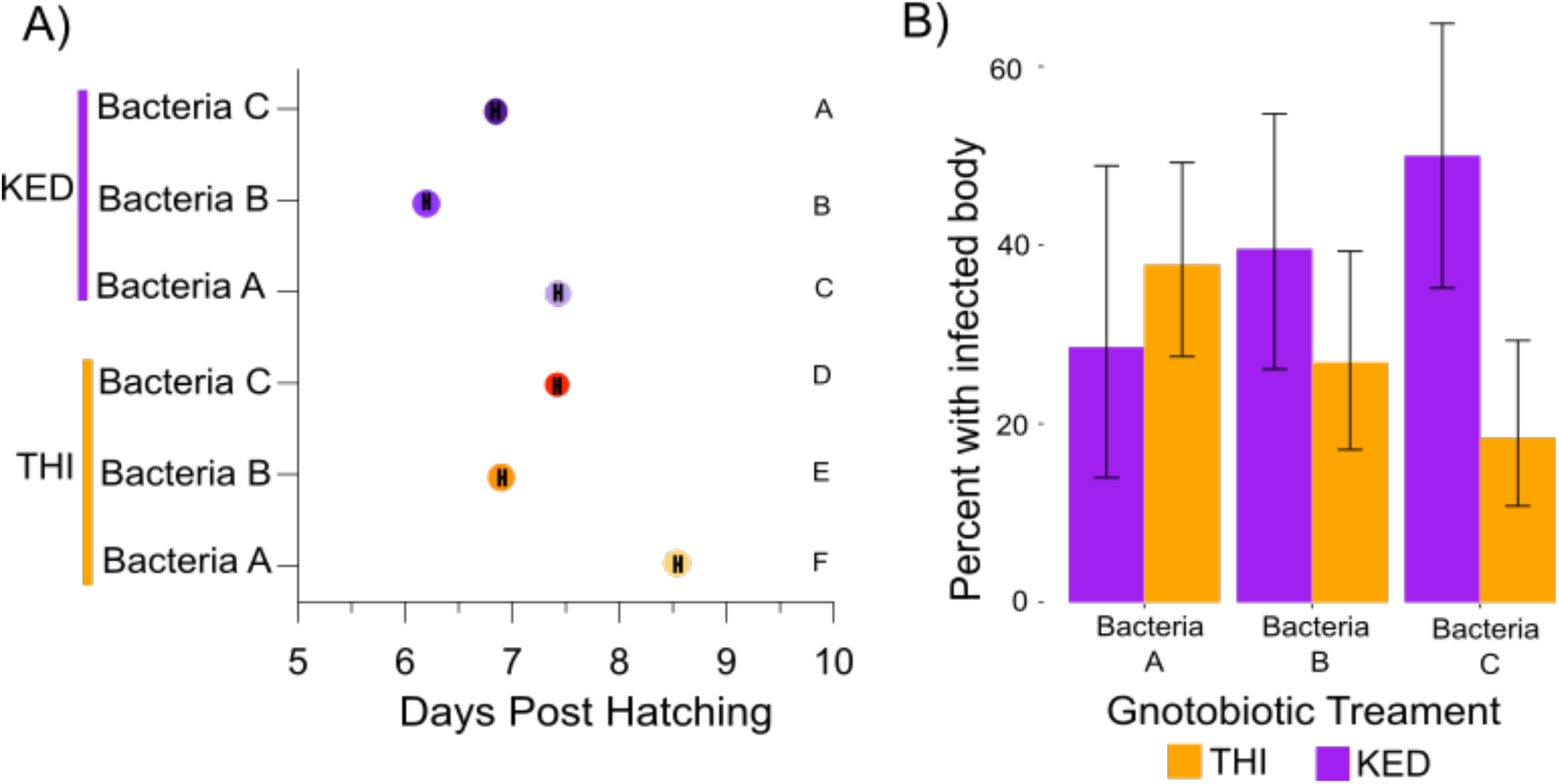
Larval exposure to different bacteria isolates alters pupation rate and ZIKV infection rates. Variation in pupation rate (A) and ZIKV infection rates (B) are shown for *Ae. aegypti* reared in the presence of single bacteria isolates in a gnotobiotic system. (A) The PD_50_ (day where approximately 50% of the larvae have pupated) is shown for two distinct genotypes of *Ae. aegypti* [7] KED and THI reared in the presence of each bacterial isolate (Bacteria A, Bacteria B, and Bacteria C). Statistical significance of differences between PD_50_ was determined by multiple comparisons with two-way-ANOVA and Tukey’s test (between bacterial treatments in each mosquito genotype) and Sidak’s test (between bacterial treatments within the same mosquito genotype) and is designated by the letters. Error bars are shown within each point. (B) The proportion of KED and THI lines of *Ae. aegypti* with a ZIKV positive body 7 days post-infection following larval rearing in single bacteria isolates, Bacteria A, Bacteria B, and Bacteria C. Error bars represent the 95% confidence intervals of the proportions. Data were analyzed by a two-way ANOVA on a binomial logistic regression (bacteria isolate: p-value = 0.614, mosquito genotype: p-value = 0.372, bacterial isolate x mosquito genotype: p-value = 0.010).

To determine if larval development in the presence of different single isolates influences ZIKV infection rates in a mosquito genotype dependent manner, the THI and KED line were exposed to ZIKV following larval development in Bacteria A, Bacteria B, or Bacteria C. Seven days post exposure to ZIKV, infection rates were determined by detection of viral RNA by RT-PCR. Infection rates ranged from 20-50% and was not dependent on the bacteria isolate (p-value = 0.614) or the mosquito genotype (p-value = 0.372), but was dependent on the interaction between the bacteria isolate and the mosquito genotype (p-value = 0.010) (Figure 1B).

### Complex microbiomes

To establish if the effect of the interaction between mosquito genotype and bacteria isolate on Zika virus infection can be extended to complex microbiomes collected from larvae in the field, whole microbiomes from the larvae pools collected in Thiés, Senegal were introduced to both genotypes of *Ae. aegypti.* We first characterized the bacterial communities by 16S amplicon sequencing. Out of 80 individual larvae sequenced, a total of 87 OTUs were found which represent 30 genera after filtering for low abundance OTUs and OTUs present in the negative controls. Rarefaction curves (Supplemental Figure 1) show that sufficient sequencing depth was achieved. Principal component analysis (PCA) was performed on a Bray-Curtis dissimilarity matrix on the larvae from the same sites to test if the bacterial communities from each collection site that were introduced to the larvae were in fact different. The community structure differed between all five of the bacterial communities (Figure 2) (PERMANOVA, p-value = 0.001). The bacterial community from Site 1 was the most different from the other sites, and Sites 2-5 showed greater similarity.

**Figure 2:**
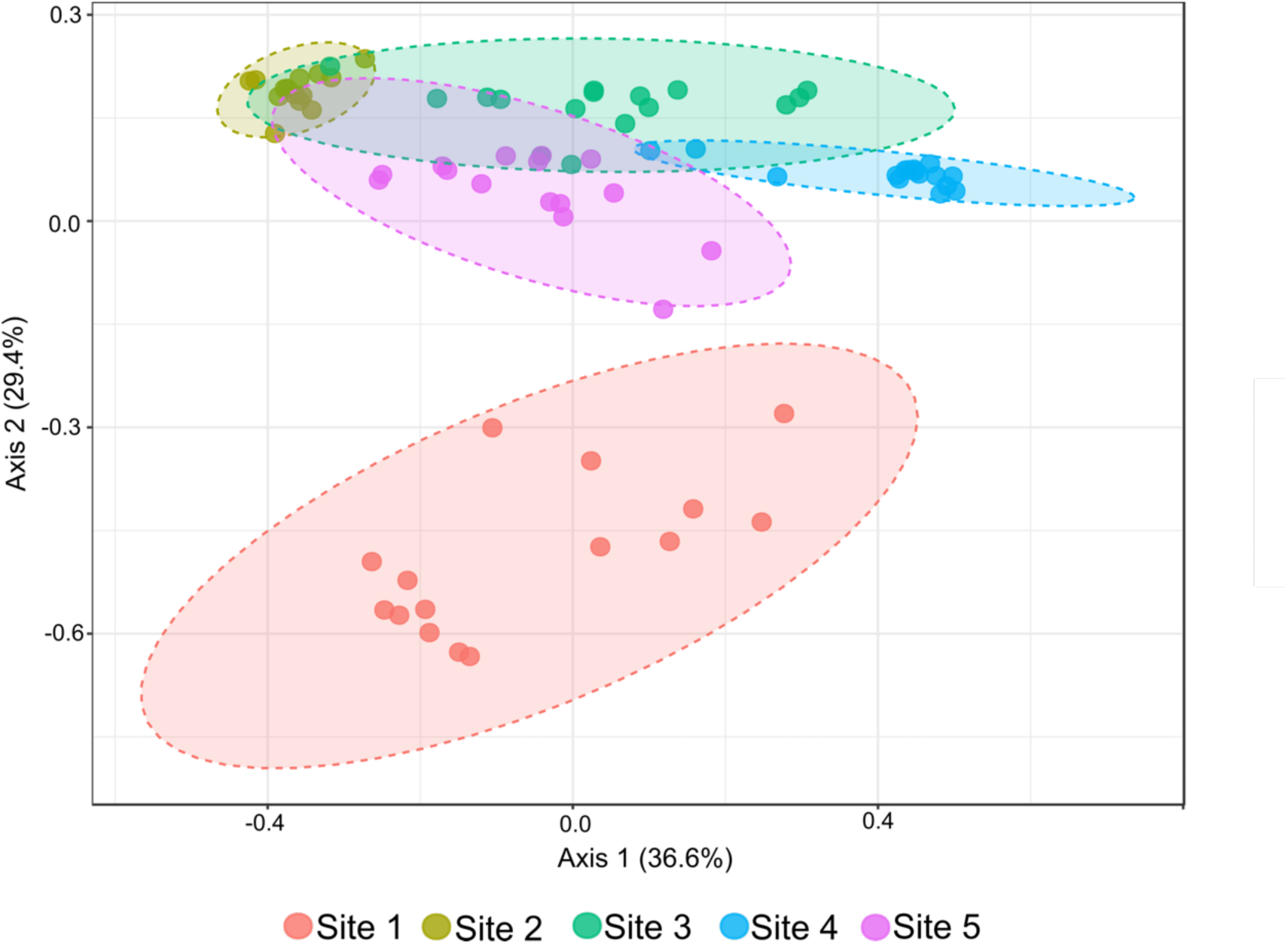
The bacterial community structure differs between the gnotobiotic larvae. Structure of bacterial communities was determined by deep sequencing the V3-V4 region of the 16S gene in individual larvae reared in a gnotobiotic system inoculated with complex microbiomes generated from larvae homogenates collected in Senegal (Site1-Site5). Bacterial structure is represented by PCoA of a Bray-Curtis dissimilarity matrix based on mean genera abundance (PERMANOVA p = 0.001).

To assess if the two genotypes of *Ae. aegypti* acquire and maintain different bacterial taxa after rearing in an identical bacterial community in a gnotobiotic system, the percent abundance of different genera was compared between genotypes and collection sites. The abundance of the top 20 most prevalent genera differed in both genotypes between the collection sites (Figure 3A). Gnotobiotic larvae harboring the bacterial community from Sites four or five are most similar between the genotypes. Gnotobiotic larvae harboring the bacterial community from Sites one, two, and three are most different between the genotypes. Abundance of specific genera was consistent between individuals from each treatment (Supplemental Figure 2). To determine if there are specific genera that are differentially abundant between genotypes across all the collection sites in the gnotobiotic system, pairwise differential abundances of each genus were compared between mosquito genotypes. Of the 30 genera identified in this study, nine genera were more abundant in the genotype from THI, and seven genera were more abundant in the genotype from KED (Figure 3B). Additionally, the overall bacterial community structure differed between mosquito genotypes when fed identical microbiomes. When all sites were analyzed together the overall bacterial community structure differed between sites and genotypes (PERMANOVA p=0.001) (Supplemental Figure 3). When each microbiome source was analyzed independently, the community structure differed between microbiomes from Sites 1, 2, and 3, but not Sites 4, or 5 (Supplemental Figure 4).

**Figure 3:**
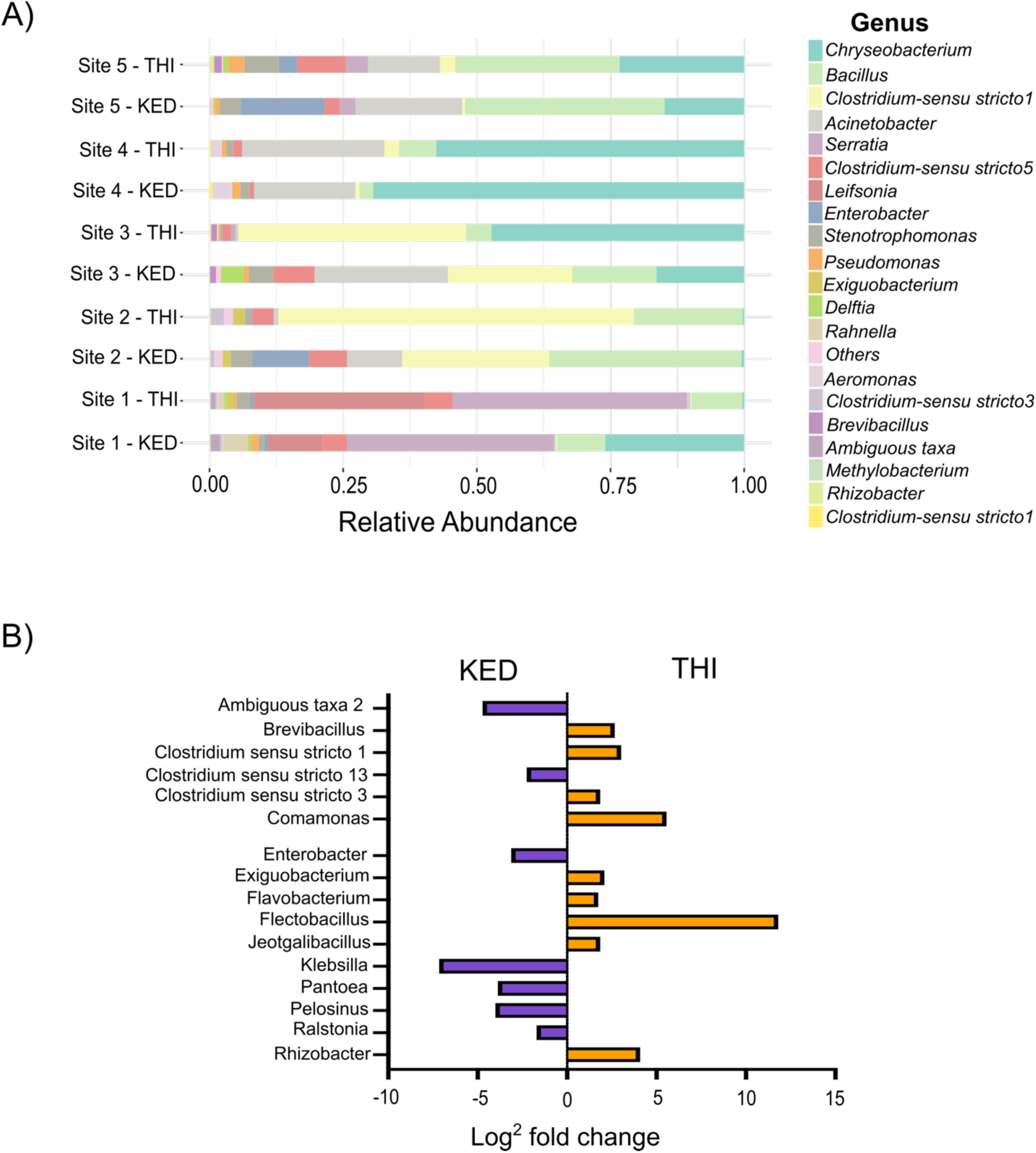
Abundance of specific taxa differs between lines of gnotobiotic larvae. (A) The percent abundance of the top 20 most abundant genera is plotted by larval treatment (Site1-5) and mosquito line (KED or THI). (B) The log_2_fold change of bacteria genera with significant pairwise differences between KED and THI is plotted. Pairwise differential analysis was performed between larvae from the KED or THI after receiving identical complex bacterial communities. The bacteria associated with negative value are the one more likely to be associated to KED and the one with positive value are the one associated with THI.

To evaluate if the mosquito genotype by larval microbiome interactions on adult susceptibility to arboviruses extends to complex microbiomes, the two different genotypes of *Ae. aegypti* were reared in preserved complex microbiomes harvested from larvae in natural breeding sites in Senegal and challenged with ZIKV. The proportion of infected individuals was determined seven days post exposure by detecting viral RNA by RT-PCR. The proportion of infected bodies varied based on the larval microbiome in which it developed and the mosquito genotype. Specifically, the proportion of the infected bodies was not dependent on the bacterial community (p-value = 0.265) or the mosquito genotype (p-value = 0.392), but was dependent on the specific pairing of larval microbiome and genotype (p-value = 0.017) (Figure 4A). To determine if this phenotype extends to dissemination titers, the number of infectious particles was enumerated in the heads seven days post infection by focus forming assay. The titer of ZIKV in the heads was dependent on the bacteria community (p-value = 0.0415), but was not dependent on the mosquito genotype (p-value = 0.057) or on the interaction between mosquito genotype by larval microbiome interaction (p-value = 0.538) (Figure 4B).

**Figure 4:**
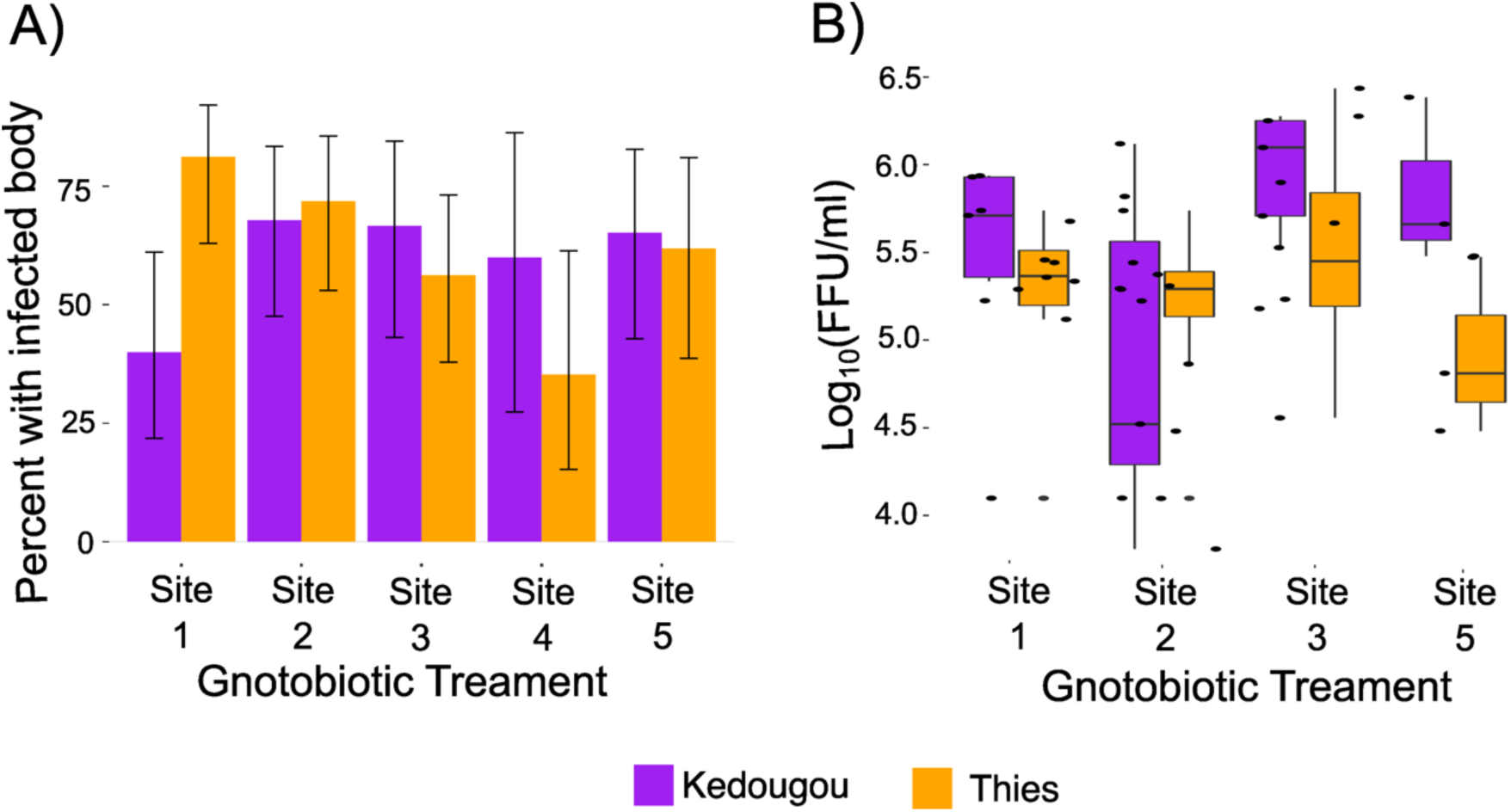
Larval development in different complex microbiomes alters ZIKV infection rates in a mosquito genotype dependent manner. (A) The proportion of blood-fed *Aedes aegypti* females from the KED and THI lines with a ZIKV-positive body 7 days post-infectious blood meal by RT-PCR following larval rearing in complex microbiomes, Site1-5. The y-axis indicates the proportion of ZIKV-infected female bodies and error bars represent the 95% confidence intervals of the proportions. Data were analyzed by binomial logistic regression as a function of bacterial treatment, mosquito genotype, and their interaction (bacterial treatment: p-value = 0.265, mosquito genotype: p-value = 0.392, but was dependent on the specific pairing of bacterial treatment x mosquito genotype: p-value = 0.017. (B) Boxplot showing the dissemination titers of infectious ZIKV particles expressed as the Log10-transformed number of focus-forming units (FFU) per ml detected in the *Ae. aegypti* head seven days post-infectious blood meal. The points represent individuals and the mean is represented by a horizontal line. The error bars represent the 95% confidence interval. Data were analyzed by Two-way ANOVA as a function of bacterial treatment, mosquito genotype, and their interaction (bacterial treatment: p-value = 0.0415, mosquito genotype: p-value = 0.057, bacterial treatment x mosquito genotype: p-value = 0.538).

## Discussion

In this study, we explored the contribution of mosquito genotype and larval microbiome in driving variation in ZIKV susceptibility. Having previously observed that adult replication of DENV was dependent on the specific bacteria the mosquito was exposed to during larval development [37], we sought to determine if the effect of larval gnotobiotic treatment on arbovirus susceptibility was mosquito genotype dependent. We found that proportion of ZIKV infected adults is dependent on the interaction between a single bacteria isolate during larval development and mosquito genotype. When reared in identical complex microbiomes in a gnotobiotic system, the mosquito genotypes differed in the abundance specific genera. Finally, we observed that the proportion of infected bodies, but not the head titers, was dependent on the specific pairing between larval microbiome and mosquito genotype. Instead, the head titers were dependent on the larval microbiome. Together these data demonstrate that different genotypes of *Ae. aegypti* interact differently with their larval microbiome and these genotype dependent interactions are important for ZIKV susceptibility.

In accordance with previously published work [37], we did not see an effect of larval development in single bacterial isolates on the proportion of infected bodies. In Dickson et al. 2017 [37], the influence of larval development in different bacteria was only observed in the amount of infectious DENV particles in the head. In the current study, we did not assay dissemination titers. If done, we might also observe differences in the amount of infectious virus outside the midgut dependent on which bacterial isolate the larvae developed in. None-the-less, we still detect an interaction between mosquito genotype and larval microbiome on the proportion of ZIKV infected bodies. Interestingly, we did observe that the amount of infectious ZIKV in the head was dependent on complex bacterial community that the larvae were exposed to in accordance with Dickson et al. 2017 [37], but no interaction between mosquito genotype and larval microbiome was detected. This might suggest that mosquito genotype by larval microbiome interactions are only important for infection rates, while the influence of the larval microbiome on the amount of disseminated virus is largely driven by the larval microbiome across mosquito genotypes.

The nutrition status and larval microbiome have previously been shown to influence mosquito fitness [41, 42] and susceptibility to arboviruses [37, 38, 43, 44]. The microbiome is composed of diverse microorganisms that colonize the mosquito’s gut, interacting with the host’s metabolic processes [38, 45, 46] and modulating its immune response [36, 47–49]. Additionally, recent research has shed light on the critical role of nutrition in determining the outcome of infection in mosquitoes. In particular, lipid metabolism plays a crucial role in the replication and dissemination of arboviruses within the mosquito’s body [50–55], while amino acids are involved in the mosquito’s interaction with the microbiome [45, 56]. Given that the mosquito’s nutritional status is strongly influences by its microbiome, and that the nutritional status of the mosquito can influence arbovirus infection, it is not surprising that we observe significant variation in the rates of ZIKV infection when mosquitoes are reared in different bacterial communities.

Additionally, there have been several mosquito genes identified that impact the microbiome composition and gut equilibrium [45, 57–60]. These genes can control the overall abundance of the microbiome or specific taxa, through their involvement in bloodmeal digestion and immune factors [57] [60]. Moreover, the mosquito microbiome has the potential to influence the expression of particular genes, which can shape the mosquito’s immune response and facilitate efficient colonization of specific microbes [59]. Although it remains uncertain which mosquito genes are responsible for the observed phenotypes, it is possible that genetic variation in genes regulating specific bacterial taxa exist between the two genotypes of *Ae. aegypti* utilized in this study.

An important finding of this study is that the different genotypes of *Ae. aegypti* larvae harbor a different abundance of specific taxa when fed identical microbiomes. Multiple studies have sought to determine if different genotypes or lines of mosquitoes have the same microbiome when maintained in the same environment. While some studies observed no differences between the microbiome between genotypes of *Ae. aegypt*i [61], other studies observe different microbiomes between lines in the same environment [62] and these changes hold up across microbially diverse environments [63]. While it is assumed that the bacteria the mosquitoes are exposed to in the same insectary is the same across lines, this is not absolute and it is very plausible that, during rearing, some larval trays could contain different microbes. By using a gnotobiotic system, we are ensuring that the different genotypes are receiving the same microbiome in a highly controlled environment. We observed differences in the abundance of specific genera between different genetic backgrounds when fed identical microbiomes. This demonstrates that the mosquito genetic background plays a role in microbiome composition. Other factors that contribute to microbiome composition are the environmentally available bacteria [46], and competition between bacteria [64]. Given that the microbiome is remodeled from larvae to adult development it is likely that that adult microbiomes are different than what is reported here and we cannot conclude that the adult microbiome is different between the two genotypes. Nevertheless, this work further highlights the importance of mosquito genotype or mosquito line in shaping the microbiome.

The results observed in this study could be dependent on the ZIKV isolate used. The isolate of ZIKV originated from Senegal and has higher infection and dissemination rates than epidemic isolates of ZIKV [65]. Perhaps we could detect an interaction between mosquito genotype and larval microbiome on dissemination titers if we had used a different isolate of ZIKV.

Even though these microbiomes originated from larvae collected in Senegal, we cannot make any conclusions of their field relevance given that we likely partially altered the composition of the microbiomes compared to larvae in the original habitat through preservation and transport. How well the microbiomes recapitulate microbiomes in nature is not relevant for this study. We simply aim to show that different complex bacterial communities can impact ZIKV infection in a mosquito genotype dependent manner and using microbiomes harvested from larvae in the field is more relevant than ad hoc mixing single bacterial isolates, even if our complex bacterial communities are not identical to those in the field. Perhaps if we seeded our gnotobiotic flasks with the microbiomes from more larvae in accordance with recent studies showing you can transplant and preserve the microbiome [66], we could make conclusions based on the origin of microbiomes.

Two of the single bacterial isolates used in this study (Bacteria A and Bacteria C) belong to the same genus, yet we observed differences in the pupation rate and infection rates between these two bacteria. The influence of these isolates on pupation rate is consistent across mosquito genotypes, but the influence of these isolates on ZIKV infection is the opposite direction across mosquito genotypes. In fact, the influence of these two isolates is likely driving the observed interaction between bacterial isolate and mosquito genotype. Perhaps genetic differences between these closely related bacteria species have variable interactions with different mosquito genotypes which are important for ZIKV infection. This system could provide a highly tractable system to investigate the mechanism of larval microbiome by mosquito interactions on arboviruses susceptibility.

Taken together these results show that different genotypes of *Ae. aegypti* interact with their environment differently to influence ZIKV infection. Future studies should expand on this work to identify how different microbiomes influence the nutrition status of the mosquito in a mosquito genotype dependent manner.

## Materials and Methods

### Bacterial Isolation

Mosquito larvae were collected in two sites (Dixième and Keur Dabo Ndione) in Thiès, Senegal from plastic drums. A pool of 3 larvae from each site was rinsed in sterile 1X PBS (Phosphate-buffered saline), incubated in 70% Ethanol for 5 minute (min), and then rinsed in sterile 1X PBS three times. Next, they were homogenized in 500 µl sterile 1X PBS and had 30% glycerol added.

A portion of the glycerol stock containing homogenized larvae was plated on Trypticase Soy Agar (rich media) and incubated for 3 days at 30 °C. Individual colonies were picked from the plates and used to inoculate 3 ml of LB media, which were shaken at 30°C until bacterial growth occurred (1 OD) and used to create new glycerol stocks of the individual isolates. DNA was extracted from each colony with the QIAGEN Dnaesy blood and tissue kit following the manufacture’s protocol. The bacterial DNA was used to amplify the entire 16*S* region by PCR [5′-AGAGTTTGATCCTGGCTCAG-3′ (forward) and 5′-AAGGAGGTGATCCAGCCGCA-3′ (reverse)] using Expand High-Fidelity Polymerase (Sigma-Aldrich). The PCR products were purified using the QIAquick PCR Purification kit (Qiagen), quantified by NanoDrop (NanoDrop Technologies Inc.), and sequenced by Sanger sequencing (Molecular Genetics Facility at University of Texas Medical Branch). The sequences were aligned and classified at the genus level using the SILVA database (www.arb-silva.de/). To standardize the amount of bacteria that would be introduced into the gnotobiotic system, aliquots of equal amounts of bacteria were made. The amount of aliquoted bacteria was then quantified by enumerating colony forming unit (CFU) for each bacterial isolate. To make the aliquots, 200 μl of each bacterial glycerol was disposed in 200 ml of LB and shaken at 30°C until bacterial growth occurred, then 50 ml was pelleted by 3000 rpm centrifugation for 15 min. The pellet was washed two times with 50 ml fresh LB broth. After the second wash, the pellet was resuspended in 50 ml of LB and aliquots were made by mixing 500 μl of resuspended bacteria an 500 μl of 50% glycerol to make 1ml aliquots. To quantify the amount of bacteria in each stock, 10 μl was taken from an aliquot and serially diluted and plated on LB plates. The number of colonies were counted and the number of CFU/ml were calculated.

### Gnotobiotic larvae

To create axenic larvae, *Aedes aegypti* eggs were collected from seventh-generation and eighth-generation laboratory colonies of Thiés (THI) and Kédougou (KED), respectively, derived from natural populations from Thiés, Senegal, and Kédougou, Senegal [7]. Genomic data for these lines exist and they represent different genotypes of *Ae. aegypti* [7]. Eggs were gently scrapped off the paper into a 50 ml falcon tube. The eggs were sterilized by incubation in 70% ethanol for 5 min, 3% bleach for 3 min, and 70% ethanol for 5 min. The eggs were then rinsed in distillated (d) sterile water three times and then they were allowed to hatch in 30 ml of d-water in a 50ml falcon tube with a 0.2 μM filter lid. Upon hatching, as a control, 10-15 axenic larvae were transferred to a sterile 25 cm^2^ tissue-culture flask containing 15ml of d-water and 50 μl of sterile fish food (1 g ground fish food flakes per 10 ml d water autoclaved for 20 minutes at 121 °C). These axenic larvae were used as an egg-sterilization control and did not develop past the 1^st^ instar larval stage in accordance with previously published work [67]. Gnotobiotic larvae were made by distributing 50 ± 5 (T-75 cm^2^ tissue-culture flasks) or 80 ± 20 (T-150 cm^2^ tissue-culture flasks) axenic larvae to sterile 75 or 150 cm^2^ tissue-culture flasks in duplicate or triplicate containing either 45 or 120 ml of d-sterile water and 1 ml of sterile fish food. For the single isolate, 5 x10 ^5^ CFUs/ml of washed-bacteria (for details go to bacteria growth section) was added to each flask. For each bacterial isolate tested, three replicate flasks were used. A total of three independent experiments was performed. To make gnotobiotic larvae with complex microbiomes, equal amounts of the glycerol stocks from each collection site (Sites 1-5) were added to each of two duplicate T-150 flasks. Data represents one experimental replicate due to availability of field material. Control and gnotobiotic larvae were maintained on 50 (T-75 flask) and 500 (T-150 flask) µl sterile fish food every other day, respectively. Bacterial isolates were chosen based on the presence of the corresponding genera in the 16S amplicon sequencing dataset.

### 16S and metagenomic analysis

To characterize the microbiome in the gnotobiotic larvae seeded with the complex microbiomes, eight individual larvae were collected from each treatment flask and transferred to a 96-well cell culture plate. The larvae were surface sterilized in 70% ethanol for 5 min and rinsed 3 times in sterile water. Next, individual larvae were transferred to 2 ml tubes containing a 5mm grinding bead and placed in the −80°C freezer until DNA was extracted. Individual mosquitoes were homogenized for 3 minutes at a 30Hz/s frequency in a TissueLyser II grinder (Qiagen). DNA extraction of individual larvae was carried out using the QIAamp DNA Kit (Qiagen, Germany) following the manufacturer’s protocol. No-mosquito controls were used for each extraction batch and included in the sequencing run.

Sequencing libraries for each isolate were generated using universal 16S rRNA V3-V4 region primers [68] in accordance with Illumina 16S rRNA metagenomic sequencing library protocols. DNA concentrations of each library were determined by Qubit and equal amounts of DNA from each barcoded library were pooled prior to sequencing. The samples were barcoded for multiplexing using Nextera XT Index Kit v2. The pooled libraries were diluted to 4 pM and run on the Illumina Miseq using a MiSeq Reagent Kit v2 (500-cycles).

To identify known bacteria, sequences were analyzed using the CLC Genomics Workbench 21.0.5 Microbial Genomics Module (CLC MGM). Reads containing nucleotides below the quality threshold of 0.05 (using the modified Richard Mott algorithm) and those with two or more unknown nucleotides or sequencing adapters were trimmed out. Reference-based Operational Taxonomic Unit (OTU) picking was performed using the SILVA SSU v132 97% database [69]. Sequences present in more than one copy but not clustered to the database were placed into de novo OTUs (97% similarity) and aligned against the reference database with an 80% similarity threshold to assign the “closest” taxonomical name where possible. Chimeras were removed from the dataset if the absolute crossover cost was three using a k-mer size of six. OTUs with a combined abundance of less than two were removed from the analysis. Low abundance OTUs were removed from the analysis if their combined abundance was below 10 or 0.1% of reads. Differential abundance analysis was performed using CLC MGM at the genus level to compare the differences between the groups using trimmed mean of M-values. Each OTU was modeled as a separate generalized linear model, where it is assumed that abundances follow a negative binomial distribution. The Wald test was used to determine significance between groups. Multiple comparisons were corrected for with Bonferroni correction. Tables of differentially abundant taxa are on the complete unfiltered data set (Supplemental Tables 2).

Abundance profiling was performed using MicrobiomeAnalyst [70, 71]. The analysis parameters were set so that OTUs had to have a count of at least 10 in 20% of the samples and above 10% inter-quantile range. Analysis was performed using actual and total sum scale abundances. Alpha diversity was measured using the observed features to identify the community richness using Chao1. Statistical significance was calculated using T-test/ANOVA. Beta diversity was calculated using the Bray-Curtis dissimilarity measure (genus level). Permutational Multivariate Analysis of Variance (PERMANOVA) analysis was used to measure effect size and significance on beta diversity for grouping variables [72]. Relative abundance analysis was done in MicrobiomeAnalyst at the level of genera.

Out of 80 individual larvae sequenced, a total of 1768 OTUs were identified. After filtering, 92 OTUs remain which represent 30 genera. Sequences from three individuals were removed from the analysis because they did not achieve enough reads, one from Site 1, 4, and 6. After removing OTUs belonging to the negative control, 87 OTUs remained. These final 87 OTUs were used for the analysis in the MicrobiomeAnalyst.

### Pupal development

To determine the rate of pupation, pupae were counted from the onset of pupation (Day 5) until pupation finished (Day 10) in the same triplicate flasks used for the adult viral challenge assays. The leftover larvae were counted and considered as total amount of individuals for the statistical analysis. For the pupal development, data from two independent experiments was used, each with three internal replicates (three replicate flasks). Graphpad was used to generate a simple logistic regression that computed the day that 50% of larvae pupated (PD_50_). An ANOVA was run on the summary statistics of the PD_50_ generated from the logistic regression to determine if the PD_50_ was dependent on the bacterial isolate, the mosquito genotype, or an interaction between the two. Multiple comparison by-two-way-ANOVAs were performed to compare the mean PD_50_ between populations (Sidak’s test), and between each of three different bacterial treatments within population (Tukey’s test) and between each bacterium in the two populations (Sidak’s test). Data are a summary of two biological experiments done each time in triplicates. The number of larvae used per experiment along with statistical information associated with each comparison are listed in Supplementary Table 1.

### Mosquito infections

Mosquito infection assays was conducted using the ZIKV DAKAR 41524 isolate received from the World Reference Center for Emerging Viruses and Arboviruses at UTMB. After pupation, pupae were transferred to a 1-pint carton box with netting and 10% sucrose solution until adult emergence. After adult emergence, 4 to 5-day-old females were sorted and transferred into a new cup with netting and deprived of sucrose solution for 24 hours and transferred to an Arthropod Containment level 2 facility (ACL-2). Females were offered an artificial blood meal for 15 minutes using the Hemotek system with de-salted pig intestine as the membrane. The infectious blood meal consisted of a 2:1 mixture of defibrinated sheep blood (Colorado Serum Company) and virus at a final concentration of 1.49 x 10^7^ focus-forming units (FFU)/ml. The blood meal was supplemented with 10 mM adenosine triphosphate (ATP). Prior to addition to the blood, sodium bicarbonate was mixed with the virus stock at 1% final concentration. Following exposure to an infectious bloodmeal, fully engorged females were sorted into 1-pint carton boxes with *ad libidum* access to 10% sucrose solution and kept in an incubator under controlled conditions (28°C, 12h:12h light: dark cycle). After 7 days of incubation, the head and body of ZIKV-exposed mosquitoes were separated to determine infection rate (the proportion of blood-fed mosquitoes with ZIKV-positive body) and dissemination titer (the amount of virus in the head tissues of ZIKV-infected mosquitoes). To determine the infection rate, female bodies were homogenized in 200 µl of a crude RNA extraction buffer (10 mM Tris HCl, 50 mM NaCl, 1.25 mM EDTA, fresh 0.35 g/L proteinase K) during two rounds of 3 minutes at a 30Hz/s frequency in a TissueLyser II grinder (Qiagen). Total RNA was converted into complementary DNA (cDNA) using M-MLV reverse transcriptase (Invitrogen) and random hexamers, the reaction was carried out as follows: 10 min at 25°C, 50 min at 37°C, and 15 min 70°C. The cDNA was amplified by PCR carried out in a 25µl reaction containing 12.5µl of 1x DreamTaq DNA polymerase (Thermo Fisher Scientific) and 10 µM of each ZIKV primer (forward: 5’-GTATGGAATGGAGATAAGGCCCA-3’, and reverse: 5’-ACCAGCACTGCCATTGATGTGC-3’). Cycling conditions were as follow, 2 min at 95°C, followed by 35 cycles of 30s at 95°C, 30s at 60°C, and 30s at 72°C with a final extension step of 7 min at 72°C. Amplicons were visualized on a 2% agarose gel. The proportion of ZIKV-infected females was analyzed by binomial logistic regression as a function of treatment, colony, and their interaction in R.

To determine the dissemination titer, the heads of females with positive ZIKV-infected bodies were titrated by focus-forming assay in Vero cells. Heads were homogenized individually in 200 µl of Vero cell media (DMEM 1X) supplemented with 2% heat inactivated fetal bovine serum (FBS) and 1X Antibiotic-Antimycotic (Life Technologies) for 3 minutes at a 30Hz/s frequency in a TissueLyser II grinder (Qiagen). Vero cells were seeded in 24-well plates and incubated for 24 hours to reach confluency. Each well was inoculated with 200µl of head homogenate in 10-fold dilutions (from 10^1^ to 10^6^) and incubated at 37°C (5% CO^2^) for 1 hour, rocking every 15 minutes. Infected cells were overlaid with α-MEM media supplemented with 1.25% carboxymethyl cellulose, 5% FBS, and 1% Pen-Strep. After three days of incubation at 37°C, infected cells were fixed with 10 % formaldehyde for at least 1 hour and cells were washed three times in 1X PBS. Approximately 500 µl of blocking solution (5% w/v non-fat powdered milk in 1X PBS) was added to each well and the plates were placed on the plate rocker for 30 minutes. The blocking solution was discarded and 200 µl of primary antibody solution (ZIKV antibody diluted 1:1000 in blocking solution) was added to each well and plates were placed on plate rocker overnight. The primary antibody solution was discarded, and plates were washed three times with 1X PBS again, and 200 µl of secondary antibody (peroxidase-labeled goat anti-mouse IgG human serum KPL-474-1806) solution (secondary antibody diluted 1:2000 in blocking solution) was added to each well. Plates were placed on plate rocker for 1 hour. The secondary antibody solution was discarded, and plates were washed three times with 1X PBS. To develop visible foci, 100µl of TrueBlue peroxidase substrate (KPL 5510-0050) was added to each well, and plates were placed on the plate rocker until foci could be seen, around 10 min. Plates were washed with deionized water and FFU was counted with the help of a light. Focus-forming units were Log_10_ transformed to represent the concentration of infectious ZIKV particles detected in *Ae. aegypti* heads. Head titer data were analyzed by two-way ANOVA as a function of bacterial treatment, mosquito genotype, and their interaction in R. Infection rate data was analyzed by a two-way ANOVA on a binomial logistic regress in R.

## Acknowledgements

We would like to thank Jiehua Zhou and Ruimei Yun of the UTMB insectary core. We would also like to thank Assyatou Gueye and Marieme Gueye for their assistance in field sampling. We would like to thank Dr. Noah Rose and Dr. Lindy McBride for sharing the mosquito colonies with us. LBD was supported by UTMB start-up funds and U01AI151801 West African Center for Emerging Infectious Diseases. AG was supported by U01AI151801 West African Center for Emerging Infectious Diseases.

## Supplemental Figures

**Supplemental Figure 1:**
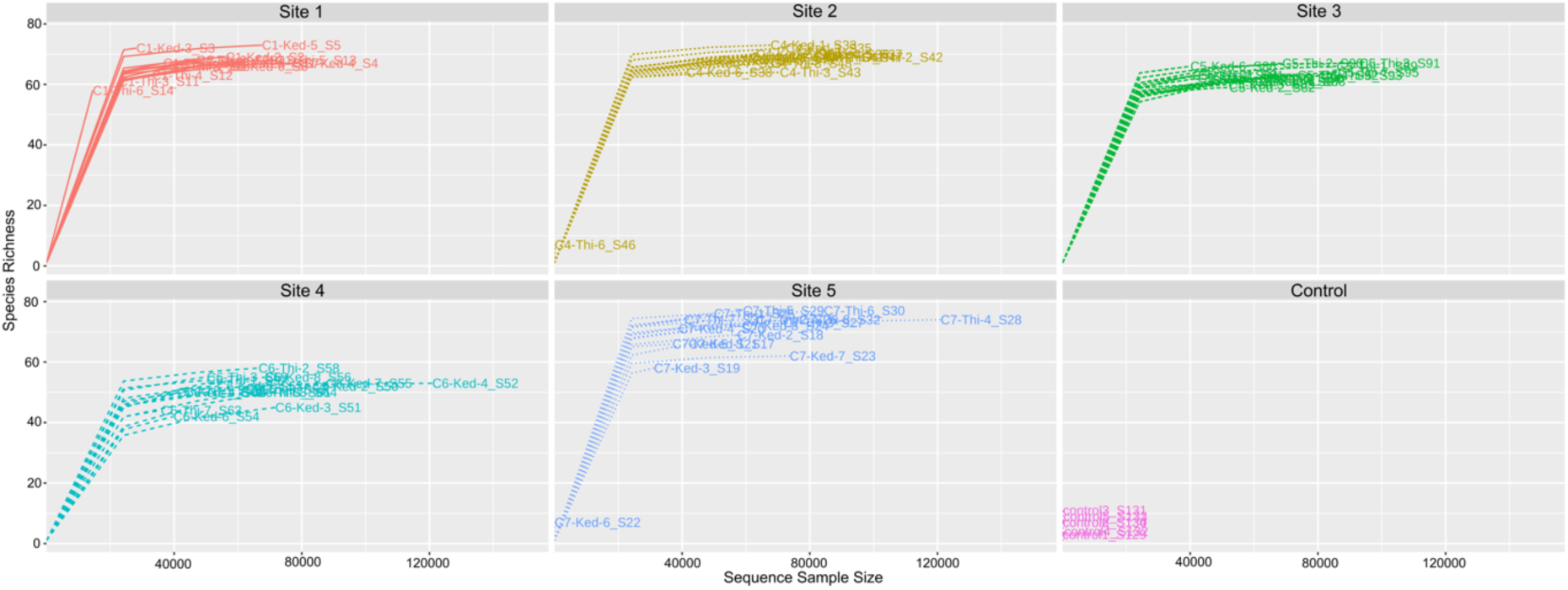
Rarefaction curves showing the sequencing depth of each library. The number of species is shown on the Y axis, and the number of sequencing reads is shown on the X axis.

**Supplemental Figure 2:**
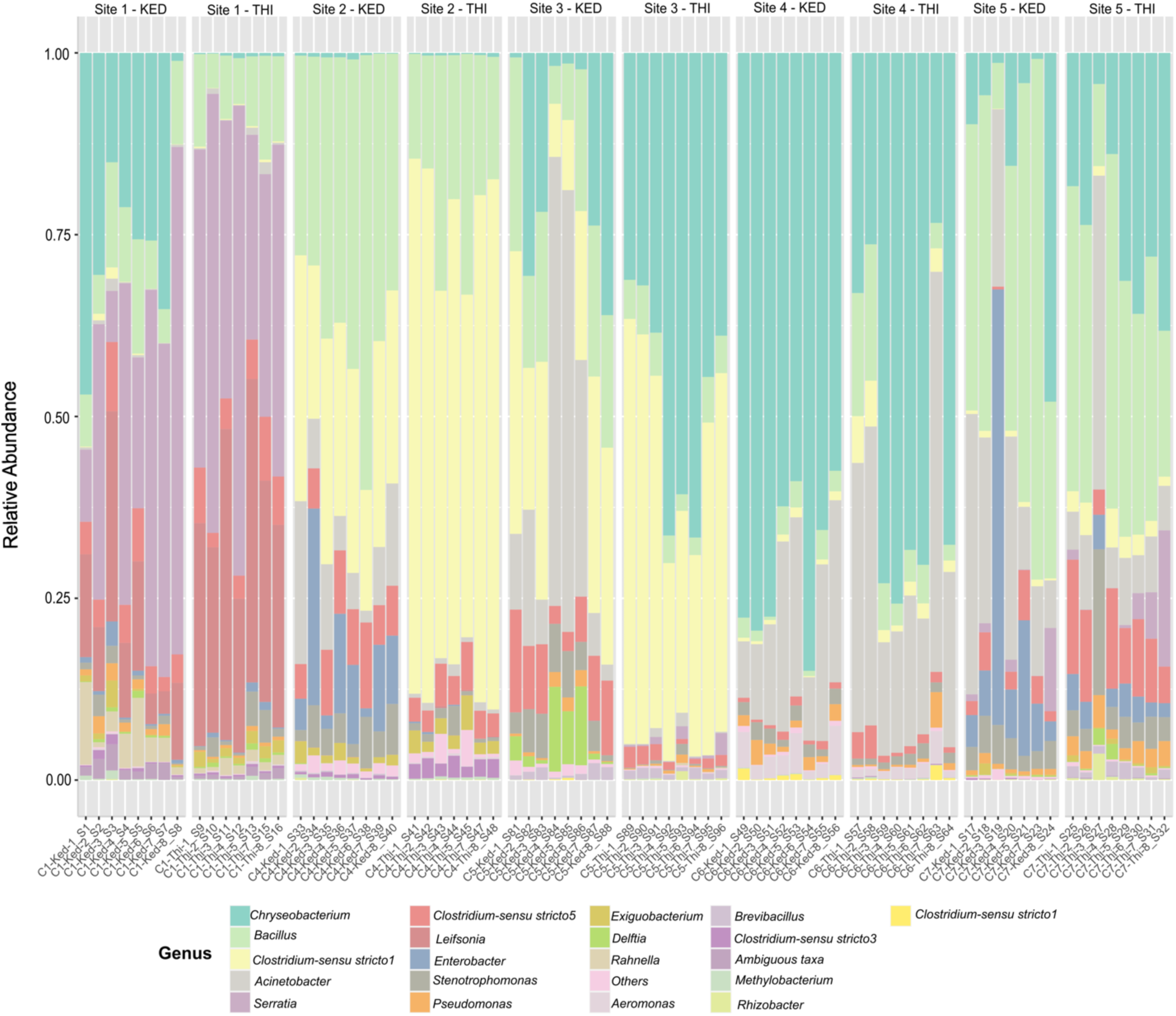
The percent abundance of the top 20 most abundant genera is plotted by larval treatment (Site1-5) and mosquito line (KED or THI) and separated out by individual.

**Supplemental Figure 3:**
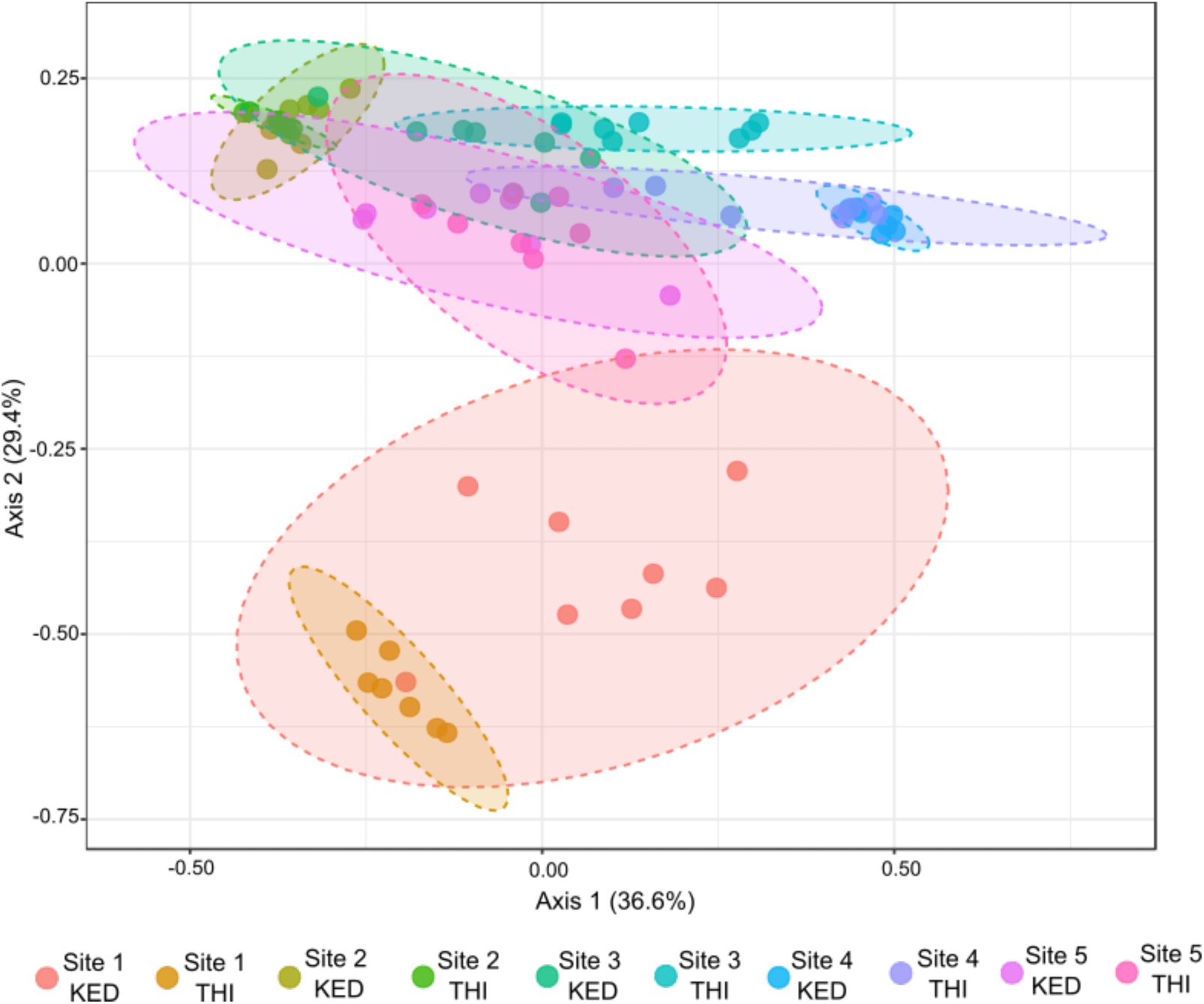
Beta diversity metrics for the *Ae. aegypti* lines in each bacterial treatment. The dissimilarities between the two different lines of *Ae. aegypti (*KED and THI) in each of the five different larval microbiomes was analyzed by principal component analysis of Bray-Curtis dissimilarity index (PERMANOVA, p = 0.001).

**Supplemental Figure 4:**
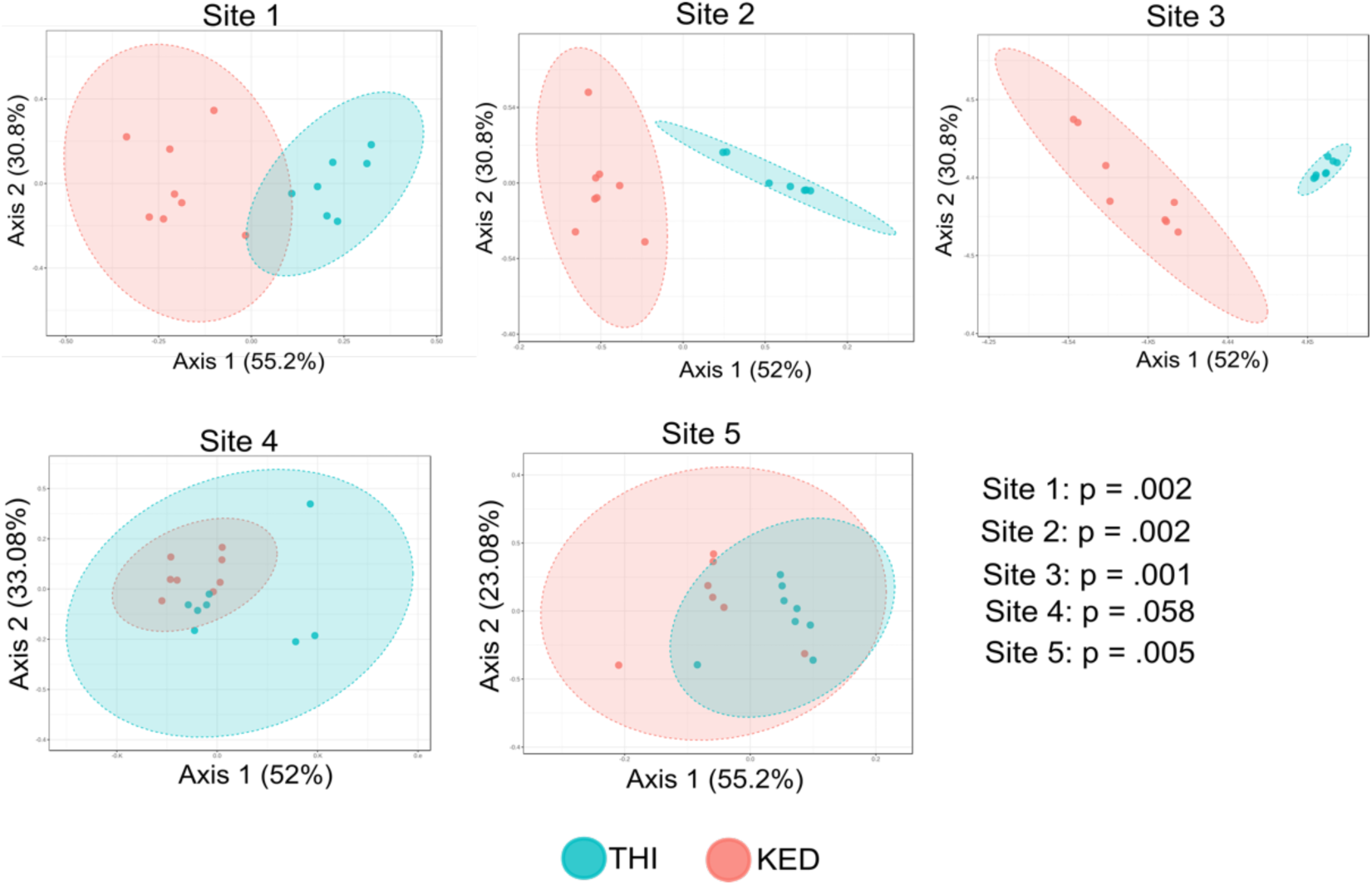
Beta diversity metrics for the *Ae. aegypti* lines in each bacterial treatment analyzed separately. The dissimilarities between the two different lines of *Ae. aegypti (*KED and THI) in each of the five different larval microbiomes was analyzed by principal component analysis of Bray-Curtis dissimilarity matrixes. P-values from individual PERMANOVA analysis are shown on the figure.

